# Population genomics of *Saccharomyces cerevisiae* human isolates: passengers, colonizers, invaders

**DOI:** 10.1101/001891

**Authors:** Carlotta De Filippo, Monica Di Paola, Irene Stefanini, Lisa Rizzetto, Luisa Berná, Matteo Ramazzotti, Leonardo Dapporto, Damariz Rivero, Ivo G. Gut, Marta Gut, Mónica Bayés, Jean-Luc Legras, Roberto Viola, Cristina Massi-Benedetti, Antonella De Luca, Luigina Romani, Paolo Lionetti, Duccio Cavalieri

## Abstract

The quest for the ecological niches of *Saccharomyces cerevisiae* ranged from wineries to oaks and more recently to the gut of *Crabro* Wasps. Here we propose the role of the human gut in shaping *S. cerevisiae* evolution, presenting the genetic structure of a previously unknown population of yeasts, associated with Crohn’s disease, providing evidence for clonal expansion within human’s gut. To understand the role of immune function in the human-yeast interaction we classified strains according to their immunomodulatory properties, discovering a set of genetically homogeneous isolates, capable of inducing anti-inflammatory signals *via* regulatory T cells proliferation, and on the contrary, a positive association between strain mosaicism and ability to elicit inflammatory, IL-17 driven, immune responses. The approach integrating genomics with immune phenotyping showed selection on genes involved in sporulation and cell wall remodeling as central for the evolution of *S. cerevisiae* Crohn’s strains from passengers to commensals to potential pathogens.

## Introduction

Despite the deep knowledge on genetic, molecular, and phenotypic traits regulating the physiology of *Saccharomyces cerevisiae*, the forces shaping its origin and evolution are still debated. The long lasting association of baker’s yeast *S. cerevisiae* with human activities (3150 b.C.) (Cavalieri et al. 2003), lead to the idea that its use in fermentation caused its domestication (Pretorius 2000). A handful of studies have examined the population biology of *S. cerevisiae* (Aa et al. 2006; Ezov et al. 2006; Fay and Benavides 2005; Legras et al. 2007; Ruderfer et al. 2006). Recently more powerful analyses based on extensive whole genome analysis of *S. cerevisiae* and *S. paradoxus* populations demonstrated that the population structure of *S. cerevisiae* consists of a few well-defined, geographically isolated lineages and many different mosaics of these lineages, supporting the idea that human influence provided the opportunity for cross-breeding and production of new combinations of pre-existing variations and then dispersed them in the environment (Liti et al. 2009). The results of genome wide analyses classified some isolates of this species with respect to the type of human activity from which they derived (Dunn et al. 2012), yet how human intervention shaped baker’s yeast structure is still unclear. The fact that divergent strains have been isolated from niches not associated with fermentation, such as oak trees, soil, and insects, suggests that *S. cerevisiae* is not domesticated generally (Borneman et al. 2011; Liti et al. 2009; Schacherer et al. 2009). Population genetics studies examining populations deriving from soil/plant niches (Ezov et al. 2006; Sampaio and Goncalves 2008; Sniegowski et al. 2002) and the interaction between fermentations and the environment (Goddard et al. 2010) provide evidence for the role of insects in local dispersal of strains, that can further serve as a seed for wine fermentation, and be clonally amplified during industrial wine production. The recent discovery of the role of wasp’s gut as a winter sanctuary and potential evolutionary niche for *S. cerevisiae* (Stefanini et al. 2012) indicated how the ecological cycle of this microorganism is just beginning to be understood. The role of the gut in *S. cerevisiae* evolution is not limited to wasps but likely extends to mammals. Humans lead inextricably mycotic lives; yeasts are ubiquitous in human related environments, leavening their bread and fermenting their wine and beer. Therefore it is not surprising that yeasts inhabit their skin, mouth, and intestinal tract. Since exposure to fungi is constant, recent studies have begun to note that fungal microflora, recently referred as *mycobiota*, is not a negligible player in host-microbe interactions. Analyses of human skin and gut mycobiota, revealed a rich community of *Ascomycetales*, such as *Candida*, *Aspergillus* and surprisingly *Saccharomyces* spp. (Findley et al. 2013; Ghannoum et al. 2010; Iliev et al. 2012; Ott et al. 2008). A few studies investigating fungal communities in Inflammatory Bowel Diseases (IBD) described a more heterogeneous mycobiota in patients with Crohn’s disease (CD) (Iliev et al. 2012) compared with healthy subjects. Further studies in mice models with DSS-induced colitis, highlighted the contribution of mycobiota to gut inflammation boost (Iliev et al. 2012), intriguingly indicating a decrease in *S. cerevisiae* and increase in *Candida* levels in inflammatory conditions. Strikingly, amongst the IBDs one of the markers proposed to discriminate CD from Ulcerative Colitis (UC) was the positivity for anti-*Saccharomyces cerevisiae* antibodies (ASCA) (McKenzie et al. 1990; Quinton et al. 1998; Sendid et al. 1998). In this work we investigate whether children with IBD are colonized by *S. cerevisiae* and what is the population structure of *S. cerevisiae* in CD patients.

## Results

### Crohn’s patients harbor a peculiar *S. cerevisiae* population evolved within the human gut

We initially used selective media to evaluate the cultivable fungal flora in feces from 34 pediatric CD patients, 27 UC patients, and 32 healthy children (HC, Supplemental Table 1). The classification according to the species was carried out by ITS1-4 region sequencing (Supplemental Tables 2-3). This analysis identified 34 isolates as *S. cerevisiae,* of these 31 strains from CD patients, 2 from UC and 2 from HC (Supplemental Fig. 1A-C). In order to understand if the fecal *S. cerevisiae* strains isolated from CD were foodborne or environmental we investigated the population structure through two genetic markers: three strain origin-mimicking sequences (Ramazzotti et al. 2012) and 12 selected microsatellite loci (Legras et al. 2007) (Fig. 1 and Supplemental Table 3), previously used to genotype and track the origin of isolates from a wide variety of sources. Both markers showed that the majority of fecal strains belonged to a single cluster, previously unknown, that we called the Human/Laboratory (H/L) cluster, since it contained also the laboratory strains S288c, W303, and Sigma1278b (Mortimer 2000) (Fig. 1A). Strikingly, strains from the same CD patient in at least 3 cases (H, P and B) were highly phylogenetically related, thus indicating clonal expansion and evolution within the host (Fig. 1 and Supplemental Fig. 1D). In particular, the fact that the four strains isolated from a single CD patient (YH1-YH4) co-clustered with strains isolated in West Africa (Fig. 1A), strongly supported the idea that host colonization and clonal amplification from *S. cerevisiae*, as strains belonging to this clade, could be isolated neither from the patient’s geographical location nor from human-related fermentation processes, thus they could only have evolved within the human gut. Those strains also showed low genetic heterozygosity (Magwene et al. 2011), probably arising from inbreeding (Supplemental Fig. 1E). Only two CD strains, YB7 and YB8 fell within the wine-European (W/E) cluster, while one UC strain fell between the West African/Crohn (WA/C) and the H/L clusters. Interestingly, the overall portrait, including the identification of outlier strains, was confirmed both by gene markers and microsatellites (Fig. 1).

**Figure 1.**
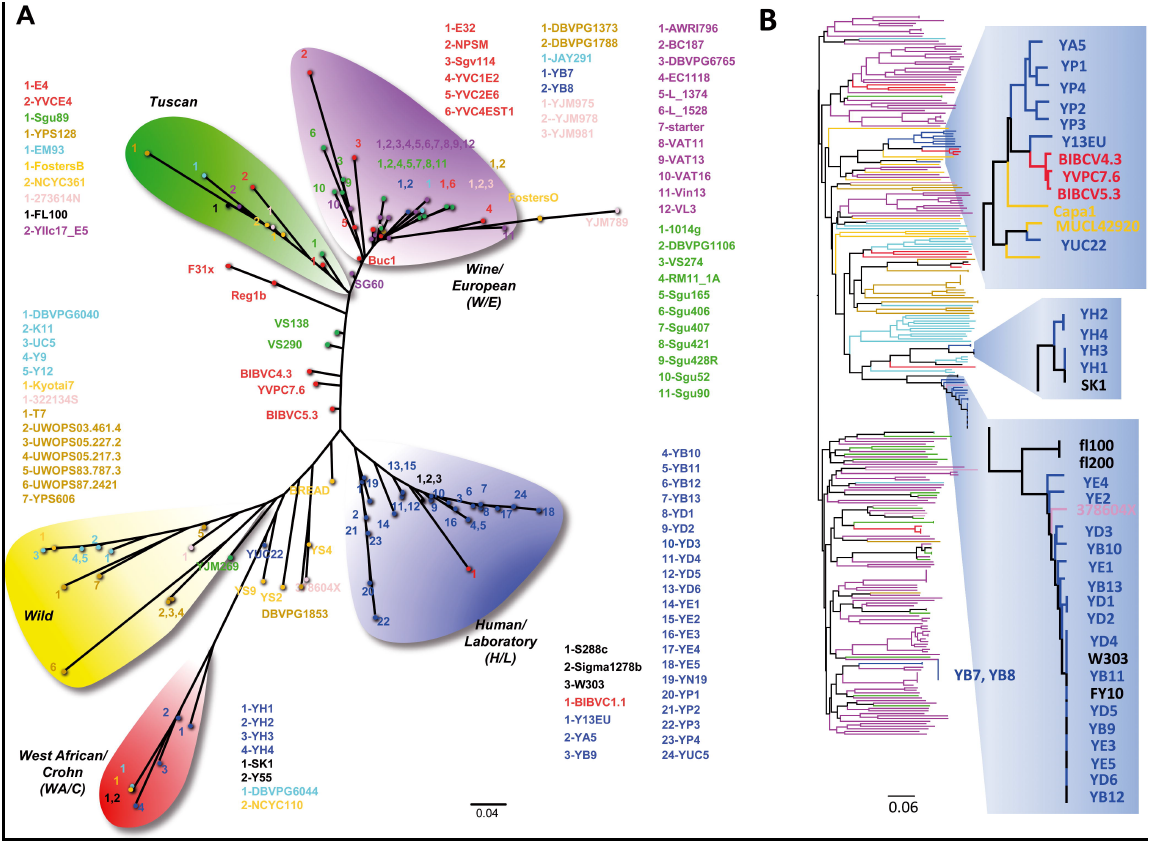
Reconstruction of *S. cerevisiae* population phylogeny. (*A*) *S. cerevisiae* strain cluster based on the *EXO5*-*IRC8*-*URN1* gene SNP sequences. (*B*) Clustering based on the genetic distances calculated with microsatellite analysis. (*A-B*) Color code: purple-wine, green-grapes, dark yellow-natural sources, yellow-bakery (bread and beer), light blue-other fermentations, pink-clinical, red-insect guts, blue-fecal isolates.

To further investigate the genomic structure of the human isolates, we completely sequenced the genomes of 31 *S. cerevisiae* selected strains and meiotic segregants from the same geographic region but from different sources. Specifically, we sequenced 19 isolates from feces, 4 from wasp guts and 9 from fruit and wines (Supplemental Material and Supplemental Table 4), representing the widest genetic variability inferred with the two sets of genetic markers. The existence of different evolutionary forces shaping the CD strain genomes was further supported by identification of a higher level of polymorphism (with respect to the S288c reference strain) in fecal isolates, compared to strains from other sources (69,038 SNPs in CD versus 40,761 SNPs on average, respectively, Supplemental Table 5). Interestingly, at the whole genome level only two fecal isolates, one CD and one UC, fell close to the laboratory strains, five strains fell within the W/E, one in the WA/C, and four fell in a separate highly different cluster together with one of the wasp strains from Chianti (Tuscany, Italy, Fig. 2A).

**Figure 2.**
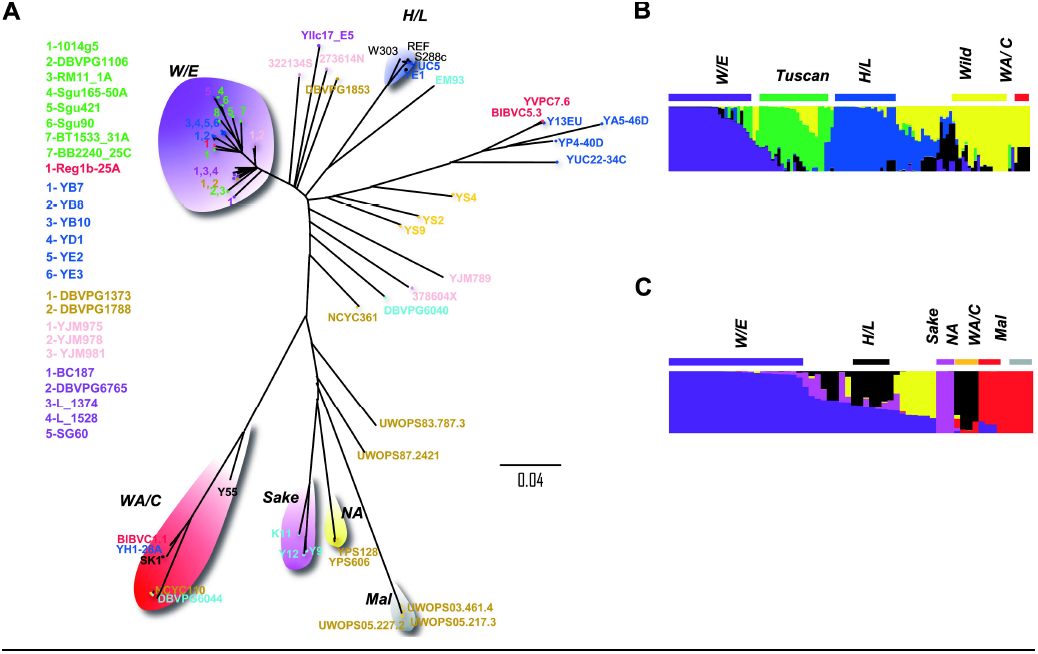
Whole-genome sequenced *S. cerevisiae* population phylogeny. (*A*) Phylogenetic tree based on the genetic distances of 21 whole genome-sequenced strains. Color code as in Fig. 1. (*B*-*C*) Strains’ most probable ancestry, inferred from Structure analysis and combined with CLUMPP, performed respectively (*B*) on the three genome-mimicking gene SNP sequences and (*C*) on whole-genome SNPs. The proportion of the most probable inferred ancestors for each strain is indicated by the colors of vertical bars. W/E, Wine/European; H/L, Human/Laboratory; WA/C, West African/Crohn; NA, North America; Mal, Malaysian.

Ancestry analysis performed on both genetic markers (Fig. 2B and C) showed that the YA5, Y13EU, and YUC22 strains and the YP series associated with different lineages from those found by microsatellite and marker gene phylogeny had a mosaic genome. The fact that strains previously associated with the H/L cluster were assigned to the W/E clade by genome wide analysis, can be attributed to widespread genome mosaicism amongst gut strains (Fig. 2C). On the contrary, the 5 strains of the CD specific clusters show a higher level of inbreeding (Supplemental Fig. 1D-E), in agreement with the results from microsatellite analysis that show that Human strains are more inbred with respect to wine and wasp strains (Fig. 1B). In contrast to the CD strains the 3 sequenced wasp strains are less inbred and fall in the 3 major distant clusters, even though they are derived from the same geographical region.

The presence of two distinct types of fecal strains can be explained by the hypothesis that different passenger *S. cerevisiae* strains can mate within the Crohn’s gut, giving rise to mosaic strains that can colonize and evolve in the human intestine as long term residents. Following sporulation and self mating within the gut, resident strains can give rise to highly homozygous strains, thus suggesting the role of the intestine as a potential generator of yeast diversity. Geography does not play a major role in shaping yeast population diversity. Rather, the different ecological niches present in the human gut, in grape fermentation and fermentation of different wine products are the main forces driving yeast evolution, that likely occurs by opportunistic blooming and clonal expansion upon exposure to environments rich in simple sugars.

### Evolution within the gut determines selective pressure in genes involved in cell wall structure and morphology

For each sequenced strain, we identified genetic losses and acquisitions with respect to the reference genome (Supplemental Fig. 2 and Supplemental Tables 6-9). We found 97 genes lost in mosaic fecal strains (Table S8) and a total of 506 new genes, 114 of which in the mosaic strains, 155 in W/E, and 85 in the WA/C strains (Supplemental Fig. 2B and Supplemental Table 9). The CD strains belonging to the W/E and WA/C clades shared 34 new genes, some of which are involved in cell morphology and potentially important for host colonization (Supplemental Table 9). Assessment of mean mutation frequency (Joseph and Hall 2004) showed that the genes involved in steroid biosynthesis and sucrose degradation were subjected to a higher selective pressure in the gut environment (Supplemental Table 10). It is worth noting that all the results from genome analysis of the gut strains converge on genes involved in biogenesis and assembling of the fungal cell wall, the structure recognized by the human immune system (Romani 2011), thus leading us to surmise different immune reactivity of these isolates.

### Sporulation ability of gut strains correlates with ASCA serum level of CD patients and disease activity

It is well known, specifically in CD, that a loss of tolerance towards commensal yeasts is expressed by serological levels of ASCA against yeast cell wall mannans (McKenzie et al. 1990; Quinton et al. 1998; Sendid et al. 1998). In this study the clinical status and ASCA serum level of IBD patients correlated with the presence/absence of yeast strains in feces (*P*=1.42×10^−6^, χ2 test). Nevertheless, no significant correlation between ASCA positive (ASCA+) CD patients and the most abundant fungal species *per se* could be found, indicating that ASCA production is not promoted by a specific yeast species, but rather by strain-specific antigenic properties present in several species, as previously observed (McKenzie et al. 1990). We further explored the hypothesis by evaluating the presence of virulence-related traits in *S. cerevisiae* isolates (Supplemental Fig. 3). Amongst the most widely established pathogenicity- and inflammation-associated traits, we studied invasiveness, resistance to temperature and oxidative stress, and sporulation (Supplemental Table 11) (Diezmann and Dietrich 2009; McCusker et al. 1994). We found significant associations between fecal strain invasiveness and spore production (*P*=0.001, Fisher’s exact test, FET), invasiveness and resistance to oxidative stress (*P*=0.01, FET), and sporulation rate and pseudohyphal formation (*P*=0.022, FET). Strikingly, all the *S. cerevisiae* strains showing medium/high sporulation capability were isolated from ASCA+ CD patients, while strains unable to sporulate were isolated from ASCA negative (ASCA-) CD patients (*P*<0.0001, FET), highlighting the pivotal role of the yeast sporal form in eliciting the host immune response.

We then investigated whether sporulation could influence yeast adaptation to the host, and therefore affect the population structure. Microsatellite-based clustering of CD isolates reflected the strain’s ability to sporulate (Fig. 1B). Fecal strains with poor sporulation ability (none or <10%) clustered with the laboratory strains S288c, Sigma1278b and W303.

Interestingly, yeast strains isolated from an ASCA+ CD patient (H) with high inflammation indexes and a UC patient (UC22) showed the highest percentage of sporulation (>30%, i.e. YH1, YH2, YH3 and YH4). These strains, belonging to the WA/C cluster, bore a highly mutated *CDA1* locus (Supplemental Fig. 4), encoding a protein involved in the biosynthesis of chitosan, a component of the yeast spore wall, further supporting the correlation between the sporulation ability of fecal strains and ASCA+. Furthermore, the sporulation proficient strains showed the *RME1* sequence, one of the two master regulators of the sporulation process, (Deutschbauer and Davis 2005), identical to that of the high sporulator SK1 (Supplemental Fig. 5 and Supplemental Table 3). While all the CD strains isolated from ASCA-patients showed the less active S288c allele.

ASCA is only one of the markers used in CD assessment. Stratifying the patients according to the levels of calprotectin, a biological marker useful for monitoring mucosal disease activity (Supplemental Table 1), a total of 18 CD patients presented high mucosal inflammation indexes, in 11/18 CD we found yeast and in 5/11 CD *S. cerevisiae.* A total of 16 CD presented low levels of calprotectin, suggesting mucosal healing; in 3/16 CD we found yeast, and in 2/3 cases *S. cerevisiae* (see Supplemental Material). All mosaic-high sporulating strains were isolated in patients with active disease. The contrary was not true since one patient (D) with active disease and high inflammation indexes had strains incapabable of sporulating, which belonged to the H/L cluster.

To assess whether the strain-specific traits observed determine differences in their immunogenicity, we measured the cytokine profiles *in vitro* on healthy human peripheral blood mononucleated cells (PBMCs), and *in vivo* on wild type and specific knock-out mice, upon challenges with IBD and HC yeasts. Despite wide intra-individual variability, the *in vitro* assay revealed IL-17A, IFN-γ (both known to afford colonization resistance), and IL-10 (the cytokine of tolerance) (Romani 2011), as the main discriminating factors among strains (Fig. 3 and Supplemental Fig. 6). Strains characterized by pure W/E ancestral lineage showed a tendency towards the induction of INF-γ-mediated responses. Strains with mosaic lineages and low sporulation efficiency induced high IL-17A production. An opposite trend was observed for the high sporulator YH1 and SK1 strains, in which the high IFN-γ mediated-inflammatory response was counterbalanced by high levels of IL-10 (Fig. 3A and B). CD patient immune responses showed greater variability compared to healthy volunteers (Supplemental Figs. 7 and 8), ranging from high IL-17A and low IFN-γ and IL-10 (YB8 and YD1) to a pure Th1-driving response (high IFN-γ and low IL-17A and IL-10, as YE5) or high IL-10 (YH1).

**Figure 3.**
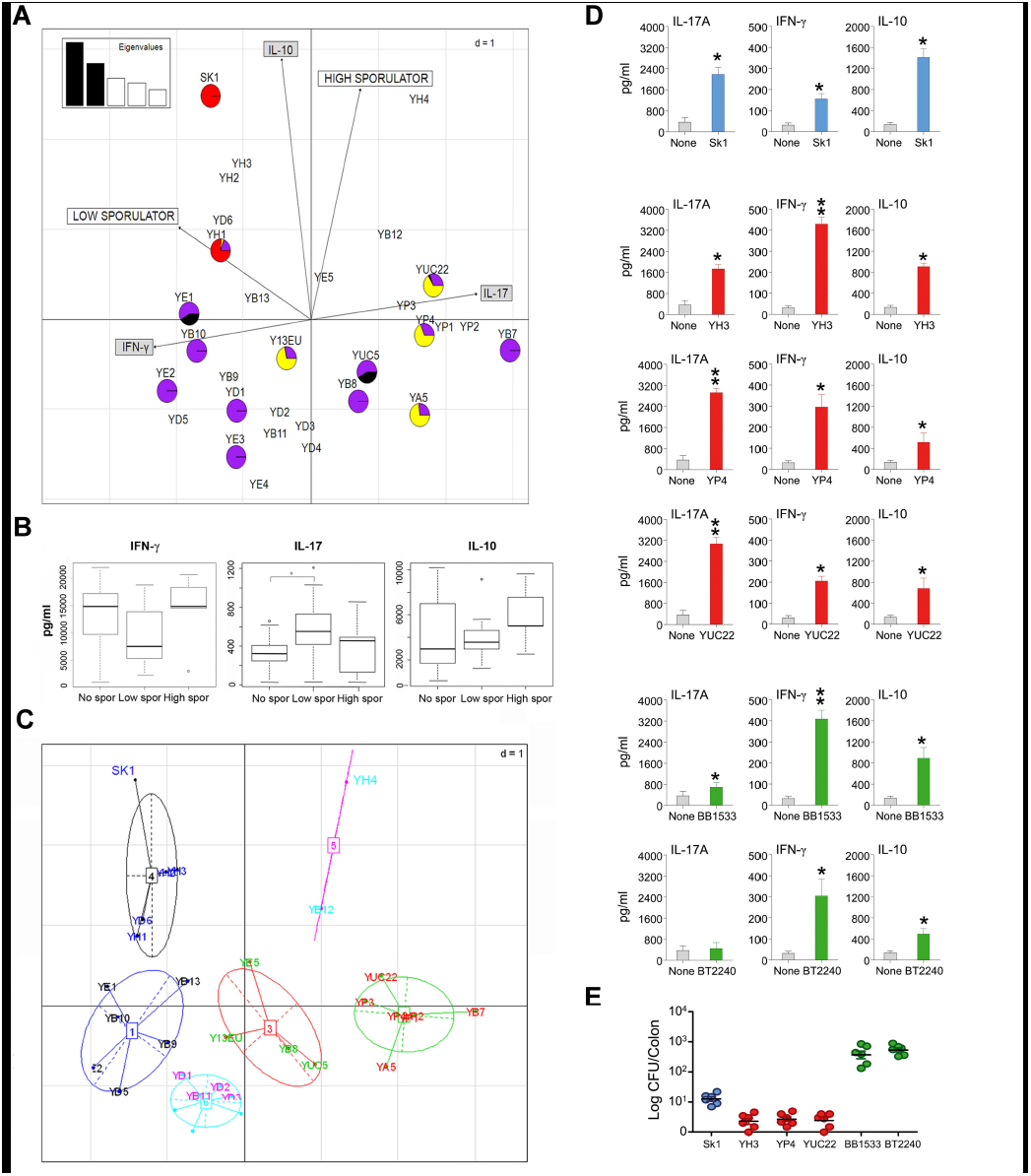
T-polarizing cytokine release in humans and mice induced by *S. cerevisiae* isolates. (*A*) First two components obtained by correspondence analysis. Cases are *S. cerevisiae* strains and variables are T-polarizing cytokine release by human PBMCs in response to the strains and the sporulation rate. Pie charts indicate the ancestor lineage proportion of each whole-genome sequenced strain (colored as in Fig. 2B). (*B*) Box plot of cytokine release (pg/ml) upon stimulation with yeast strains grouped by sporulation ability. (*C*) Visual representation of strain grouping inferred using PAM analysis of the PCA coordinates, based on the cytokine pattern. (*D*) Cytokine production at 3 days post-infection. \**P*<0.05, \*\**P*<0.01, naïve versus infected C57BL/6 mice. (*E*) Fungal growth (measured as log CFU) in the colon of C57BL/6 mice infected with selected *S. cerevisiae* isolates.

One patient (B) was sampled both in active disease and in remission (Supplemental Table 12), showing presence of *S. cerevisiae* strains in both conditions. The strains isolated in active diseases clustered amongst the W/E strains with all the markers used being sporulators and produced IL-17A. On the contrary the strains isolated in remission were unable to sporulate and produced INF-γ. This finding, again, suggests a role of sporulation ability in the inflammatory process.

Prompted by these observations, we explored the existence of alleles related to the different immune responses (Fig. 3C), leading to the discovery of 20 genes with alleles significantly correlated with: (i) low IL-10, (ii) high IL-10, or (iii) high IL-17A and low IL-10 responses, respectively. Among these, we found genes related to important mechanisms for avoiding the immune response and persisting within the host, such as genes involved in cell wall morphology and composition, hyphal formation and switching, and cell polarization (Supplemental Table 13). These 20 alleles represented precious targets for future studies on the immunomodulatory ability of *S. cerevisiae*.

In parallel, cytokine profiles in wild type C57BL/6 mice infected with mosaics (YH3, YUC22 or YP4) or pure W/E lineage (BT2240, BB1533) strains reflected the immune phenotype elicited in healthy human PBMCs (Fig. 3D and Supplemental Fig. 9). Yeast isolates differed in their ability to grow and colonize the colon at 3 days post inoculation (Fig. 3E). *S. cerevisiae* colonization was associated with inflammatory influx of the colon and positive mucin staining, particularly with mosaic isolates (Supplemental Fig. 10). To define the functional role, if any, of these cytokines in yeast colonization and/or infection, we comparatively evaluated parameters of inflammatory pathology in the colon of C57BL/6 wild-type (WT) or cytokine-deficient mice inoculated with the different isolates. Interestingly, inflammation was drastically reduced in IL-17A-deficient mice inoculated with mosaic strains, but promoted in IFN-γ-deficient mice inoculated with non mosaic isolates and in all infected IL-10-deficient mice (Supplemental Fig. 10), indicating IFN-γ and IL-10 responses are protective. Thus, *S. cerevisiae* isolates contributed differently to local epithelial activity and inflammation in the colon.

Overall, the observations strengthen the remarkable correlation between the specific genetic make-up, ASCA production, and sporulation rate. Mosaic strains were associated with ASCA+ CD patients and sporulation, and H/L or pure ancestors with ASCA- CD patients (Supplemental Fig. 11 and Supplemental Table 12), supporting the hypothesis that ASCA production is promoted by the antigens of strains with specific genetic features.

## Discussion

Here we provide evidence of a human gut adapted lineage of *S. cerevisiae*. The observation of clonality for the CD strains argues convergent evolution and selection for traits related to cell wall composition and sporulation, supplying a fertile habitat for prolonged stays and colonization. The higher frequency of *S. cerevisiae* in CD patients is likely the result of a higher permeability in the intestines of CD. The genomic analysis demonstrated a higher level of mosaicism in CD *S. cerevisiae* strains, suggesting that the human gut possibly serve as an accelerator for yeast evolution. The genetic, immunologic, and phenotypic results converge to indicate a set of genes related to cell wall composition and sporulation as associated with strain-specific differences in the cytokine pattern induced in CD and with the ASCA marker, thus reflecting the yeast’s ability to induce different inflammatory responses. These findings highlight the need to consider the interplay between fungal cell wall and gut immune function in determining mycobiota composition and evolution. From this perspective, pathogenic traits probably result from rapid convergent evolution and adaptation to different ecological niches present within the host, leading to differential yeast strain immunogenicity *via* differential exposure of cell wall antigens (Gow and Hube 2012; Rizzetto et al. 2013; Romani 2011). The opposite immunomodulatory ability of sporulating-mosaic strains versus non sporulating can be explained by two opposite evolutionary paths. In the first path, purifying selection favors strains that can evade the immune system and do not induce inflammation, in the second path, since spores are inflammatory and induce Th-17 responses (Rizzetto et al. 2010) selection within the gut favored non sporulating pure lineage strains. These non-sporulating pure strains evolve in the gut to give tolerogenic responses, and are potentially alleviating in IBD patients, but could themselves become dangerous by escaping immunosurveilance. Sporulation could have evolved as a strategy for *S. cerevisiae* to be disseminated within the environment, therefore sporulators have a potential advantage in surviving the gut of a normal host (Coluccio et al. 2008; Rizzetto et al. 2010). This advantage turns into a danger when the intestine of the host is weaker, the mucus layer is damaged, or its immune system impaired, in these cases, high sporulating strains are a potential cause of disease and inflammation. The role of this microorganism as pathogenic or commensal is thus determined from strain rather than species specific traits. From an evolutionary perspective, different environments, represented by healthy and disease conditions, could ultimately affect the ability of fungal strains to sustain or dampen chronic inflammation, and of humans to serve as reservoirs and evolutionary niches for the most useful and friendly of our food borne microorganisms, *S. cerevisiae*. We have to rethink profoundly the concept of yeast domestication, as being driven not only by selection on traits important for fermentation of fermented beverages, but also from the potential of certain fungal strains to modulate immune function in the human intestine. The evidence presented in this study bears significantly on the understanding of the role has the mycobiota on evolution of immune function in health and disease, showing how the definition of a strain as pathogenic-commensal is deeply intertwined with the status of the gut-immune barrier. The molecular mechanisms used by yeast to colonize the host, as a harmless commensal are a continuum with the strategies used for evading immune surveillance and potentially turn a friend into a threat.

## Methods

### Enrolment of patients and healthy subjects

A total of 93 pediatric subjects were enrolled at the Meyer Children’s Hospital (Florence, Italy). All individuals were Caucasian and aged range from 4-19 years (Supplemental Table 1). For CD patients, disease activity was scored using the Pediatric Crohn’s Disease Activity Index (PCDAI, see Supplemental Material). For UC patients, a similar index was used to measure disease activity, the Pediatric Ulcerative Colitis Activity Index (PUCAI, Supporting Text). Inflammatory activity was assessed in all IBD patients through clinical parameters, such as erythrocyte sedimentation rate (ESR), C Reactive Protein (CRP) and fecal calprotectin, a useful marker of mucosal inflammation (Aomatsu et al. 2011). Anti-*Saccharomyces cerevisiae* Antibodies (ASCA) IgA and IgG levels were evaluated for every IBD patient (Supplemental Material and Supplemental Table 1). All enrolled individuals were made aware of the nature of the experiment, and all gave written informed consent in accordance with the sampling protocol approved by the Ethical Committees of the Meyer Children’s Hospital and the Azienda Ospedaliera Universitaria Careggi, Florence, Italy (Ref. n. 87/10).

### Isolation and identification of yeast species from fecal samples

Feces were collected from all pediatric subjects. A 1ml feces aliquot was plated on Yeast Extract-Peptone-Dextrose (YPD) agar medium supplemented with chloramphenicol (1mg/ml) and incubated for 2-3 days at 28° C. Yeast genomic DNA was extracted as previously described (Hoffman and Winston 1987). Strains were identified by sequencing of the ribosomal Internal Transcribed Spacer (ITS) region, using ITS1 and ITS4 primers (Supplemental Table 3), as previously described (Sebastiani et al. 2002).

### Statistical analysis and correlation among clinical parameters and yeast isolates

To evaluate variables that might influence the presence of yeast in fecal samples, such as clinical parameters, sex, age, and location of the inflammation and treatment (Supplemental Table 1), we performed logistic regression through the use of automated model selection. To observe correlations between yeast isolates from IBD and ASCA, we used the Wald test. Chi Square statistics were performed to associate significant correlations with the variables mentioned above. Yates correction was applied in the case of expected frequencies less than 5.

### Sanger sequencing of fecal *S. cerevisiae* strains

Three genome-mimicking genes, *EXO5*, *IRC8* and *URN1*, capable of recapitulating the whole genome phylogeny, were sequenced (Ramazzotti et al. 2012) by using the primers listed in Supplemental Table 3. The *RME1* gene was sequenced using the primers listed in Supplemental Table 3. Sequences of samples analyzed (Supplemental Table 4), submitted to GenBank (ID: KF261601-KF261724), were compared with *S. cerevisiae* pre-edited sequences downloaded from the Sanger institute website. The collection included strains isolated from a wide variety of sources, such as grapes, vineyards, wasps, and different types of fermentation, as well as other clinical strains (isolated in concomitance with other fungal infections). The Neighbor-joining tree was built on distances calculated with the Kimura-two parameter method (Fig. 1A).

### Population genetic analysis

Population ancestries were estimated by using the model-based program Structure (Pritchard et al. 2000). K=5 and K=6 were chosen as the most representative of the population structures for the three genome-mimicking gene sequences. The results of 10 independent Structure chains were combined with CLUMPP. Population structure was also evaluated using principal component analysis (PCA) and Differential Analysis of Principal Components (DAPC) using the R adegenet package. The groups’ inter- and intra- genetic distances, calculated with the Kimura two parameter model, were statistically tested using the Mann-Whitney U test and Levene test.

Inbreeding was evaluated by considering the *S. cerevisiae* population investigated as a whole (Supplemental Fig. 1E). FIS was evaluated for non-haploidized strains using the R adegenet package and compared with values inferred from Fstat. Intra- and inter- group FIS variance was evaluated in strains gathered according to origin and clade partnership as in Fig. 1. The Wilcoxon rank sum test and Brown-Forsyth test (Levene test on medians) were calculated using the R package stats and car, respectively. FIS among strains isolated from the same CD patient (YB, YD, YE, YH, YP strains series) were calculated (Supplemental Fig. 1D and E).

### Microsatellites analysis

A set of 229 strains was characterized for their allelic variation at 12 microsatellites loci according to Legras et al.(Legras et al. 2007). The chord distance Dc matrix was calculated for each strain with a laboratory-made program. The phylogenetic tree was obtained from the distance matrices with Neighbor of the Phylip 3.67 package, and drawn up using MEGA4.0 (Tamura 2011). The tree was rooted using the midpoint method. To assess the assignment of each fecal isolate to a specific origin, Instruct (Gao et al. 2007) was used to evaluate the number of populations that can be observed in this set of strains. The results of 10 independent chains were combined with CLUMPP, and the consensus file was illustrated with Distruct (Fig. 1B).

### Whole genome analysis

A set of strains, including isolates from paediatric IBD and HC faecal samples (N=19), wasp guts (N=4), grapes and wines (N=6), as well as an ancestor (EM93) of the S288c reference strain (Table S5) were completely sequenced. Genome Sequencing NGS runs were performed on a Genome Analyzer II (GA II), and base-called with Illumina RTA (Real-Time Analysis) Version 1.8.70.0.

The purity of the signal from each cluster was examined over the first 25 cycles and chastity = Highest_Intensity / (Highest_Intensity + Next_Highest_Intensity) was calculated for each cycle. To remove the least reliable data from the analysis, reads were filtered according to chastity> 0.6, for all but one of the first 25 bases. If there were two bases, the read was subsequently removed.

### Illumina quality control and SNP calling

Illumina reads were subjected to quality control (filtering and trimming) using QC Toolkit (http://59.163.192.90:8080/ngsqctoolkit/). Paired reads were filtered with parameters: −1 70, (cutOffReadLen4HQ) and -s 20 (cutOffQualScore). As a result, reads with a PHRED quality score less than 20 for more than 30% of their length were discarded. Moreover, reads were trimmed at the 3’end for bases having a PHRED quality score less than 30. Paired reads were mapped to the reference genome (*Saccharomyces cerevisiae* S288c, NCBI ID=559292, using Burrows Wheeler Aligner (BWA http://bio-bwa.sourceforge.net/). The Genome Analysis Toolkit (GATK) (Patel and Jain 2012) was used for base quality score recalibration, Indel realignment, duplicate removal, and to perform SNP and Indel (insertion or deletion) discovery (McKenna et al. 2010).

### Variant imposition

The coding sequences of *S. cerevisiae* strains were created using variant imposition in order to create uniform sets of genes for further analysis. Briefly, this newly developed technique inserts variants (SNPs and Indels) produced by GATK and specific for each strain into the coding sequences of the reference strain (Supplemental Material and Supplemental Table 5).

### Phylogenetic analysis

Phylogenetic analysis was carried out with Phylip (DePristo et al. 2011), using the pipeline previously developed (Ramazzotti et al. 2012). The aligned/concatenated SNPs sequences were used to compute distances with the Kimura two parameter model (Kimura) (Felsenstein 1989) using Phylip dnadist and then clustered with the Neighbour-joining method (Saitou and Nei 1987), using Phylip Neighbour.

### Gene loss prediction

For each strain, the alignments (BWA) of reads versus the reference genome were analyzed. Genes of the reference genome with coverage equal to 0 in at least one base were determined for each strain analyzed. These genes were apparently non-functional, broken, or lacking evident similarity with the reference, so we considered them as possibly lost, also considering that only a minority of genes (around 10%) had some positions (<5) with a coverage equal to 0. Technical errors could have affected this result, especially for genes belonging to regions with low complexity or repetitive sequences that could be difficult to align. Given their positions, these false positive genes should appear in the majority of the strains analyzed and could be either false positives or acquisitions of the reference genome (Supplemental Table 6).

### Discovery of new genes

For each strain, reads that did not map with the reference genome (see above) were *de novo* assembled using AbySS (Simpson et al. 2009) with *k* = 64. Contigs with a length of less than 200 bp were discarded. Gene prediction was performed using Augustus (Stanke et al. 2008), filtering out predicted genes with a length less than 100 bp. The translated sequences were then searched for similarity versus non-redundant protein database of NCBI using BLASTP (Altschul et al. 1990), with an e-value cut-off of 1e-10 (Supplemental Table 9).

### Phylogenetic analysis

Phylogenetic analysis was carried out with Phylip (Saitou and Nei 1987), using the pipeline previously developed (Ramazzotti et al. 2012). The aligned/concatenated SNPs sequences were used to compute distances with the Kimura two parameter model (Kimura) (Felsenstein 1989) using Phylip dnadist and then clustered with the Neighbor-joining method (Saitou and Nei 1987), using Phylip Neighbor (Figs. 1 and 2).

### Mutation frequency (sBEF analysis)

For each genomically sequenced strain the mutation frequency of every gene was calculated as the number of SNPs/the bp length of the gene (as in the S288c reference, Supplemental Table 10). These estimations were used as values in counter-intuitive pathway analysis. The higher the mutation frequency score, the lower the possibility that the gene function is maintained, finally mimicking gene deletion and affecting the relative pathway. These gene lists and their relative mutation frequency were analyzed with an Eu.Gene Analyzer 5.1 (Cavalieri et al. 2007), using FET. This approach describes each pathway as up- or down-regulated, considering the relative number of genes being highly or normally mutated annotated to that pathway and the result of the FET, false discovery rate-corrected, for the same pathway. In our analysis, we included yeast pathways from the KEGG (Kanehisa et al. 2006), Reactome (Joshi-Tope et al. 2005), and YOUNG (lists of target genes for known transcription factors) databases (Pritchard et al. 2000).

### Phenotypical characterization of yeast strains

*S. cerevisiae* isolates were tested for virulence related traits. We evaluated the ability of strains to rapidly adapt to environmental changes and acquire different phenotypic conformations in order to assess the potential ability of yeast cells to invade human tissues (Supplemental Fig. 3 and Supplemental Table 11). FET was performed in order to evaluate significant correlations between different virulence-related traits.

### PBMCs preparation, fungal challenge and cytokine assays

PBMCs were isolated from fresh blood obtained from 5 pediatric CD patients and 6 healthy donors, as a control. The experimental plan was approved by the local Ethical Committee of the Meyer and Careggi hospitals, and informed consent was obtained from all CD patients and healthy donors. Separation and stimulation were performed as previously described (Netea et al. 2006). All stimulations were carried out by challenging PBMCs with live fungi at 10^6^ cells/ml concentration. After 24 hr or 7 days of incubation, supernatants were collected and stored at −20°C until assayed by mean of cytokine detection. Human Milliplex^®^ assay for Tumor TNF-α, IFN-γ, IL-1β, IL-6, IL-10 and IL-17A production was performed according to the manufacturer’s instructions using Luminex technology (Fig. 3 and Supplemental Figs. 6, 8 and 9).

### Statistical analysis of human immune response data

To find possible correlations between a specific immune response and the ancestral lineage of the strains and the ability to sporulate, we carried out PCA (ade4 R package) using strains as cases and the cytokine release by human PBMCs upon stimulation with the strains as variables. Additionally, we investigated the relationship between the immune response and the strain’s sporulation ability, using the percentage of sporulation after 5 days in SPOIV medium as a factor.

We explored the existence of alleles in the yeast genomes related to different immune responses. We at first carried out Phylogenetic Analysis by Maximum likelihood (PAM) analysis on the PCA coordinates to group the strains on the basis of the immune response they elicit.

We then analyzed the concatenated genomic SNP sequences of the strain tested for the immune response, superimposing the partnership on the previously identified immunological group as phenotypes using Strat software. Alleles identified as significantly concordant to the population structure were further screened for their uniqueness in the immunological groups.

### Mice infection, histology and cytokine detection

To assess the susceptibility of C57BL/6 mice to gastrointestinal colonization and/or infection with different *S. cerevisiae* isolates, C57BL/6 female mice (6-8 weeks old) were maintained under specific pathogen-free conditions at the Animal Facility of University of Perugia (Perugia, Italy). Homozygous Il17a-/-, Ifng-/- and Il10-/- mice, on a C57BL/6 background, were bred under specific pathogen-free conditions (Fig. 3D and E and Supplemental Fig. 10). Experiments were performed according to the Italian Approved Animal Welfare Assurance A 245/2011-B.

For gastrointestinal infection, 10^8^ yeast cells were injected intragastrically. Mice were monitored for fungal growth at 3 and 7 days after intragastric inoculation in different organs, such as the esophagus, stomach, ileum, and colon, and for dissemination, liver and kidneys. Colony forming units (log10 CFU) were assessed per organ (± SEM), by serially diluting homogenates on Sabouraud dextrose agar plates and incubating at 35°C for 48 h. Cytokine content was assessed by enzyme-linked immunosorbent assays (R&D Systems) on colon homogenates.

For histology, paraffin-embedded tissue sections (3-4 µm) were stained with periodic acid-Schiff (PAS) reagent. Sections of frozen tissue, cut at 4 µm, were fixed for 60 s in methanol. Photographs were taken using the Olympus BX51 microscope at 10 or 40 (insets) × objective (Supplemental Fig. 10).

The two-tailed Student’s t test was used to compare the significance of differences between groups. A value of P<0.05 was considered significant. The data reported are representative of 4 independent experiments, with similar results. The *in vivo* groups consisted of 6 mice/group.

## Data access

All sequences generated in this study have been deposited in the Gene Bank (ID KF261601-KF261724), ENA (ID: ERP002541) and Array Express (ID: E-SYBR-8).

## Acknowledgements

This project was supported by the FP7 Integrative Project SYBARIS (Grant Agreement 242220), Ministero dell’Istruzione, dell’Università e della Ricerca, Italy Grant PRIN 2007-N352CP_001, University of Florence, Fondazione E. Mach and AMICI onlus. The authors thank M. Santos for kindly providing the Barriada wine strains, M.G. Netea and M.G. Torcia for their suggestions on immunological experiments, F. Vaggi, C. Martinez and V. Frankell for critically reading the article and C. Romualdi for critical reading and discussion on the statistical analyses of the data.

## Authors contributions

C.D.F. and M.D.P. contributed equally to this work.

P.L. and D.C. created and managed the project.

P.L., C.D.F., M.D.P. and D.C. provided and interpreted clinical data.

M.D.P. and C.D.F. performed isolation, identification and phenotypical characterization of yeast.

I.S. performed population genetic analysis.

D.C., I.G.G., M.G. and M.B. led and performed the whole genome analysis. L.B., M.R. performed assembly, SNP calling, gene loss prediction and discovered new genes analyses.

J.L.L. and I.S. performed microsatellites analysis.

M.R. and L.D. performed statistical analysis.

D.R. performed separation of yeast meiotic segregants.

L.Ri., M.D.P. and C.D.F. performed fungal challenge on human PBMCs and cytokine assays.

L.Ro., C.M.B. and A.D.L. performed fungal infection in mice and cytokine assays.

All authors contributed to write the paper.

D.C. designed the work led, supervised and integrated the different expertise and analyses.

## Disclosure declaration

The authors declare no conflict of interest.

Supplemental material is available for this article.

